# Evaluation of Transfer Learning Methods for Detecting Alzheimer’s Disease with Brain MRI

**DOI:** 10.1101/2022.08.23.505030

**Authors:** Nikhil J. Dhinagar, Sophia I. Thomopoulos, Priya Rajagopalan, Dimitris Stripelis, Jose Luis Ambite, Greg Ver Steeg, Paul M. Thompson

## Abstract

Deep neural networks show great promise for classifying brain diseases and making prognostic assessments based on neuroimaging data, but large, labeled training datasets are often required to achieve high predictive accuracy. Here we evaluated a range of *transfer learning* or pre-training strategies to create useful MRI representations for downstream tasks that lack large amounts of training data, such as Alzheimer’s disease (AD) classification. To test our proposed pre-training strategies, we analyzed 4,098 3D T1-weighted brain MRI scans from the Alzheimer’s Disease Neuroimaging Initiative (ADNI) cohort and independently validated with an out-of-distribution test set of 600 scans from the Open Access Series of Imaging Studies (OASIS3) cohort for detecting AD. First, we trained 3D and 2D convolutional neural network (CNN) architectures. We tested combinations of multiple pre-training strategies based on (1) supervised, (2) contrastive learning, and (3) self-supervised learning - using pre-training data within versus outside the MRI domain. In our experiments, the 3D CNN pre-trained with contrastive learning provided the best overall results - when fine-tuned on T1-weighted scans for AD classification - outperformed the baseline by 2.8% when trained with all of the training data from ADNI. We also show test performance as a function of the training dataset size and the chosen pre-training method. Transfer learning offered significant benefits in low data regimes, with a performance boost of 7.7%. When the pre-trained model was used for AD classification, we were able to visualize an improved clustering of test subjects’ diagnostic groups, as illustrated via a uniform manifold approximation (UMAP) projection of the high-dimensional model embedding space. Further, saliency maps indicate the additional activation regions in the brain scan using pre-training, that then maximally contributed towards the final prediction score.

## 1. INTRODUCTION

Around 55 million people suffer from dementia globally, and Alzheimer’s disease (AD) contributes to 60-70% of these cases, according to the World Health Organization (WHO).^1^ Although AD results from a progressive, abnormal accumulation of amyloid and tau proteins in the brain, this process is only measurable using highly invasive techniques such as PET and CSF sampling from the spinal cord. An MRI-based method to detect AD, even prior to more invasive testing, would be valuable, based on the distributed pattern of brain atrophy that is evident on brain MRI. AD leads to subtle and diffuse changes in anatomical features of the brain starting with the parahippocampal area and the entorhinal cortex and spreading to the rest of the cortical gray matter in a stereotypical sequence.^2^ Structural magnetic resonance imaging (MRI) based on standard anatomical T1-weighted (T1-w) is routinely used and accessible worldwide and does not subject the patient to ionizing radiation. In recent times with an increase in the available training data, deep learning and machine learning methods have shown great promise for various neuroimaging tasks such as brain age prediction,^3^ diagnostic classification^4^ and disease subtyping and staging,^5^ based on detecting profiles of brain atrophy with T1-w MRI. In 2019, Wen et al.^6^ published a comprehensive review of over 30 papers that used convolutional neural networks (CNNs) for AD classification from anatomical MRI. They found that over half of the surveyed papers may have suffered from non-independence of the training and test data, but in general they found that T1-w scans were more effective for AD detection using data driven approaches, compared to other types of MRI, such as diffusion imaging.

Traditionally, learning methods relied on large, annotated training datasets that were directly relevant to the downstream tasks of segmentation, diagnostic classification, or predictive modeling. In 2010, the launch of the ImageNet-1K dataset^7^ enabled computer vision researchers to ‘pre-train’ their models on a publicly available dataset with over a million images from 1,000 annotated classes. Pre-training - also known as *transfer learning -* defines the initial state of a model’s trainable parameters, and in some cases, some of the network weights are optimized on a prior task and then ‘frozen’, with others being allowed to be trained on the new task. This pre-training strategy can improve performance on downstream tasks, while also reducing convergence time. With the development of deeper CNNs, transfer learning on ImageNet provided a way to avoid overfitting issues in problem domains with less training data, such as medical diagnosis, where publicly available patient datasets are often limited to hundreds rather than millions of labeled examples. In 2019, M. Raghu et al.^8^ pretrained deep networks on ImageNet for medical imaging tasks, showing the benefits of feature re-use, with some dependence on other factors such as the size of the network and nuances of the fine-tuning stage. Azizi et al.^9^ also showed benefits of multiple levels of “pre-training” on both unlabeled and labeled image data. Contrastive learning, a type of self-supervised learning introduced in recent years, uses pairwise data transforms to learn high-level features without requiring additional labeling of the unlabeled data.^10^

Researchers have investigated numerous deep learning approaches for AD classification from brain scans; Martinez-Murcia et al. obtained a balanced accuracy of 0.777 on T1-w MRI from ADNI^11^ with an approach based on convolutional autoencoders. Folego et al.^12^ used a VGG-based CNN to obtain an AUC of 0.680 on MRIs from ADNI, AIBL, OASIS and other datasets, and, as noted earlier, Bin Lu et al.^13^ noted the positive effect of transfer learning on downstream neuroimaging tasks after using sex classification as the initial task. In ^6^ the authors distinguished two types of transfer learning - ImageNet-based and autoencoder (AE)-based. Valliani and Soni^14^ found that using an ImageNet pre-trained ResNet CNN improves AD diagnosis when used together with data augmentation, but they only used midline axial slices from the overall 3D MRI. One remarkable recent study^13^ created an MRI-based AD diagnostic classifier using deep transfer learning by pooling MRI data from over 217 sites and (85,721 scans from 50,876 participants). They achieved 91.3% accuracy in leave-sites-out cross-validation on the Alzheimer’s Disease Neuroimaging Initiative (ADNI) dataset, and 94.2%/87.9% accuracy for direct tests on two unseen independent datasets (AIBL/OASIS).

### Contributions

In this paper, we benchmark the effect of pre-training on diagnostic classification of AD, for a range of deep learning network architectures, datasets, and pre-training scenarios. The key contributions are:

- We tested different pre-training strategies, to learn useful feature representations for the AD classification task from brain MRI.
- We compared different deep learning architectures including 3D and 2D convolutional neural networks (CNNs), for detecting AD.
- We tested our strategies on multiple test sets to evaluate generalization across data from independent sites.
- Salience mapping methods to enhance model interpretation and better understand the features used.

## 2. DATA

### 2.1 Datasets and Data Preparation

We analyzed publicly available data from an overall cohort of 4,098 scans/1,188 subjects from the Alzheimer’s Disease Neuroimaging Initiative^15^ (ADNI), i.e., ADNI1, ADNI2 and ADNI GO, which are different phases of the ADNI project. Data from ADNI was stratified by subject ID into 3 non-overlapping sets: a training set of 2,577 scans/747 subjects, a validation set of 302 scans/84 subjects, and a test set of 1,219 scans/359 subjects. We used data from the Open Access Series of Imaging Studies, phase 3^16^ (OASIS3), as the out-of-distribution dataset to independently validate our proposed pre-training strategies. The demographics and age distributions for these splits are provided in **Table 1**. The diagnosis of dementia from ADNI and OASIS3 will be referred to as AD in this paper, although since these classifications were made, there has been some evolution and debate over how AD is best operationally defined.^17^ For one of our pre-training strategies, we also analyzed 3D T1-w brain MRI from 10,445 healthy control subjects from the UK Biobank^18^ dataset (UKBB) for supervised pre-training. The UKBB cohort was split into training (N = 7,937, mean age = 62.6(7.5) years, range: 44.5 – 80.7, 4,195F (53%)), validation (N = 418, mean age = 63.1(6.1), range: 48.9 – 75.2, 202F (48%)), testing (N = 2,090, mean age = 62.6(7.4), range: 45.4–79.7, 1,090F (52%)). For one of the pre-training strategies assessed in this paper, we initialized our model with training weights from,^19^ from a model that was trained originally on 10,420 scans/7,764 subjects aggregated from 13 sites (the ‘Big Healthy Brains’, or BHB dataset, mean age = 32(19.0) years).

**Table 1.** Brain MRI datasets analyzed in this paper, from the Alzheimer’s Disease Neuroimaging Initiative (ADNI), and the Open Access Series of Imaging Studies, phase 3 (OASIS3).

|  | ADNI | OASIS3 |
| --- | --- | --- |
| Age Range (years) | 55.2 – 95.8 | 43.5-97.0 |
| Age Average (SD) | 75.7 (6.8) | 70.6 (9.4) |
| Female Participants | 556 (46.8%) | 341 (56.8%) |
| Participants with Alzheimer’s disease | 1,704 (41.5%) | 183 (30.5%) |
| Mini-Mental State Exam(MMSE) Score:<br>Mean (range) | 26.1 (0.0 – 30.0) | 27.5 (7.0-30.0) |
| MRI scanner field strength | 2,566 3T/1,532 1.5T | 3T/1.5T |
| Number of scans | 4,098 | 600 |
| Number of subjects | 1,188 | 600 |

### 2.2 T1-weighted Image Pre-processing

The T1-w brain MRI volumes were pre-processed using a sequence of steps,^4 20^ including nonparametric intensity normalization (N4 bias field correction),^21^ ‘skull stripping’ for brain extraction, linear registration to a template with a 9 degrees of freedom transformation, and isometric voxel resampling to 2-mm resolution. The pre-processed T1-w images were of size 91×109×91. T1-w images were then z-scored to standardize the voxel intensities input to the models.

## 3. METHODS

### 3.1 Random Forest Classifier using Radiomics Features

As a baseline, we used a machine learning pipeline with radiomics features which are classical descriptors based on the image intensity histogram, classical texture-based features such as gray-level co-occurrence matrices (GLCM), and other classical image processing measures using the PyRadiomics^22^ toolbox. The top extracted radiomics features from the T1-w scans were then used to train a random forest classifier.

### 3.2 CNN Architectures

In this paper, we trained and tested 2D and 3D convolutional neural networks (CNN).

#### 3.2.1 3D DenseNet 121 CNN

The 2D version of the DenseNet^23^ architecture was originally proposed as an updated version of the popular ResNet^24^ with many more connections between subsequent layers to achieve a better performance but with fewer parameters than comparable networks. This architecture has been shown to be effective for other neuroimaging applications, including brain age prediction.^25^ The 3D CNN architecture used in this work is based on a 3D variant of the DenseNet121 architecture, as proposed in.^19^

#### 3.2.2 2D ResNet18 CNN

We used a 2D ResNet18,^24^ a standard ImageNet architecture, as a 2D CNN pre-trained with weights from the ImageNet dataset. The feature maps of the CNN’s backbone are aggregated by means of a max-pooling operation before the classifier block. Even though the DenseNet backbone is deeper than that of the ResNet, they both have approximately 11 million parameters.

### 3.3 Pre-training Strategy

Pre-training defines the initial state of the neural network - with the goal of improving performance on the downstream task. Our first pre-training strategy was a traditional supervised learning approach that uses labeled training data. We tested two different types of supervised pre-training - *intra-domain*, where we pre-trained the 3D CNN on T1-w scans from the UKBB cohort to predict the sex of the subjects, as advocated in,^13^ and *inter-domain*, where we used weights from pre-training on the benchmark ImageNet-1K dataset.^7^ Secondly, we tested a self-supervised intra-domain pre-training approach based on the contrastive learning based SimCLR framework proposed by Chen et al..^10^ SimCLR is based on minimizing a normalized temperature-scaled cross entropy loss by maximizing the agreement between 2 transformed versions of an image. In this work, we create random crops of the T1-w scans as the choice of the transformation. Lastly, we use pre-training weights for our 3D CNN provided by the authors in,^19^ which was generated using a contrastive learning framework that incorporated the chronological age of the subject’s scan as continuous proxy metadata. The weights were obtained by training on 10,420 scans from a multi-site cohort (the ‘Big Healthy Brains’, or BHB dataset).

### 3.4 Model Training

The 3D CNN model layers were initialized based on the Kaiming initialization scheme.^26^ A random search was conducted for hyperparameter optimization for our experiments including: learning rate [2e-3 to 1e-5], learning rate schedulers [scale on plateau, cosine annealing]. Fine-tuning experiments for the 3D CNN were performed end-to-end on the models including all the batch normalization layers. For the pre-training task on UKBB, the 3D CNN was trained for 50 epochs with the Adam with weight decay optimizer with a learning rate of 0.0006, weight decay of 0.0004 and a learning rate scheduler. For all the other experiments, the models were trained until there was no improvement in the validation loss for 20 consecutive epochs. The 3D CNNs used a batch size of 4 and the 2D CNN used a batch size of 1. The 2D and 3D CNNs were trained from scratch using the Adam with weight decay optimizer, an initial learning rate of 0.0006, which was reduced to 0.0001 while fine-tuning a pre-trained model. During fine-tuning, all the models minimized the binary cross-entropy loss function.

### 3.5 Model Evaluation

All models were evaluated using the receiver-operator characteristic curve-area under the curve (ROC-AUC). We also computed metrics that included the accuracy, precision, recall, and the F1-score that were calculated on the ROC curve using a threshold obtained with the Youden’s index.^27^ The model performances are presented as an average over multiple random seeds and their standard deviations. The trained models were evaluated on data from multiple sites including a test set from ADNI and an out-of-distribution test set from OASIS3.

### 3.6 Model Interpretation

We also used the Uniform Manifold Approximation (UMAP) approach^28^ to visualize the embedding space of the model for the subjects in each of the classes. We also created saliency maps to visualize the MRI brain regions that maximally contributed to the model predictions. This was achieved by calculating the gradient of the top predicted class from the models’ final layer with respect to the input image.^29^ These gradients are visualized in the Results section.

## 4. RESULTS

Overall, the pre-training methods boosted classification accuracy, especially when using 10% to 25% of the total number of available training scans, i.e., 257 scans (220 subjects) to 644 scans (438 subjects). Specifically, the 3D CNN trained from scratch for AD classification on 10% of training T1-w MRIs achieved an average ROC-AUC of 0.776 on ADNI and the fine-tuned 3D CNN yielded 0.833 using the modified contrastive pre-training (3D CNN + CL w BHB12) and 0.828 using self-supervised pre-training (3D CNN + SSL w UKBB) strategies as shown in **Table 2**. The 3D CNN pretraining task of sex prediction on scans from UKBB achieved a very high-test ROC-AUC of 0.998 (with an accuracy of 0.979, F1 of 0.987, precision 0.975, recall of 0.981). The 3D CNN trained from scratch for AD classification on 25% of the training MRIs achieved an average ROC-AUC of 0.780 on ADNI and the fine-tuned 3D CNN yielded 0.854, 0.857 and 0.792 using the modified contrastive pre-training, self-supervised pre-training and supervised (3D CNN + SL w UKBB) pre-training strategies. Using 100% of the training data i.e.,2,577 scans (747 subjects) yielded an average ROC-AUC of 0.911 on the ADNI test using the modified contrastive pre-training strategy. The baseline random forest classifier achieved an ROC-AUC of 0.723 using radiomics features.

**Table 2.** Summary of the various model architectures we trained and evaluated on the ADNI test set (CNN: convolutional neural network, SSL: self-supervised learning, CL: contrastive learning, RF: random forest), as well as the pre-training approaches. The table shows the effect of pre-training as a function of varying % of training data samples. Pre-training contributes positively to the model performance, particularly in low training data regimes. The performance is presented as the average ROC-AUC with the standard deviation with different random seeds. The results from this table are illustrated in **Figure 1.**

| Training Data % | 5% | 10% | 25% | 35% | 50% | 100% |
| --- | --- | --- | --- | --- | --- | --- |
| **3DCNN** | 0.799 (0.026) | 0.776 (0.016) | 0.780 (0.113) | 0.880 (0.005) | 0.864 (0.014) | 0.883 (0.017) |
| **3DCNN + SSL w UKBB** | 0.795 (0.014) | <u>0.828 (0.008)</u> | <u>0.857 (0.035)</u> | 0.851 (0.023) | <u>0.884 (0.009)</u> | 0.868 (0.025) |
| **3DCNN + SL w UKBB** | 0.693 (0.006) | 0.736 (0.002) | <u>0.792 (0.006)</u> | 0.845 (0.003) | 0.849 (0.007) | 0.878 (0.002) |
| **3DCNN + CL w BHB12** | <u>0.866 (0.006)</u> | <u>0.833 (0.019)</u> | <u>0.854 (0.022)</u> | <u>0.885 (0.008)</u> | <u>0.899 (0.003)</u> | <u>0.911 (0.006)</u> |
| **2DCNN** | 0.654 (0.046) | 0.646 (0.021) | 0.718 (0.024) | 0.789 (0.024) | 0.748 (0.037) | 0.779 (0.070) |
| **2DCNN + ImageNet** | 0.766 (0.026) | 0.795 (0.064) | 0.842 (0.033) | 0.589 (0.218) | 0.658 (0.172) | 0.621 (0.183) |
| **RF + radiomics** | 0.644 (0.006) | 0.685 (0.004) | 0.675 (0.037) | 0.723 (0.008) | 0.712 (0.004) | 0.723 (0.001) |

**Figure 1.**
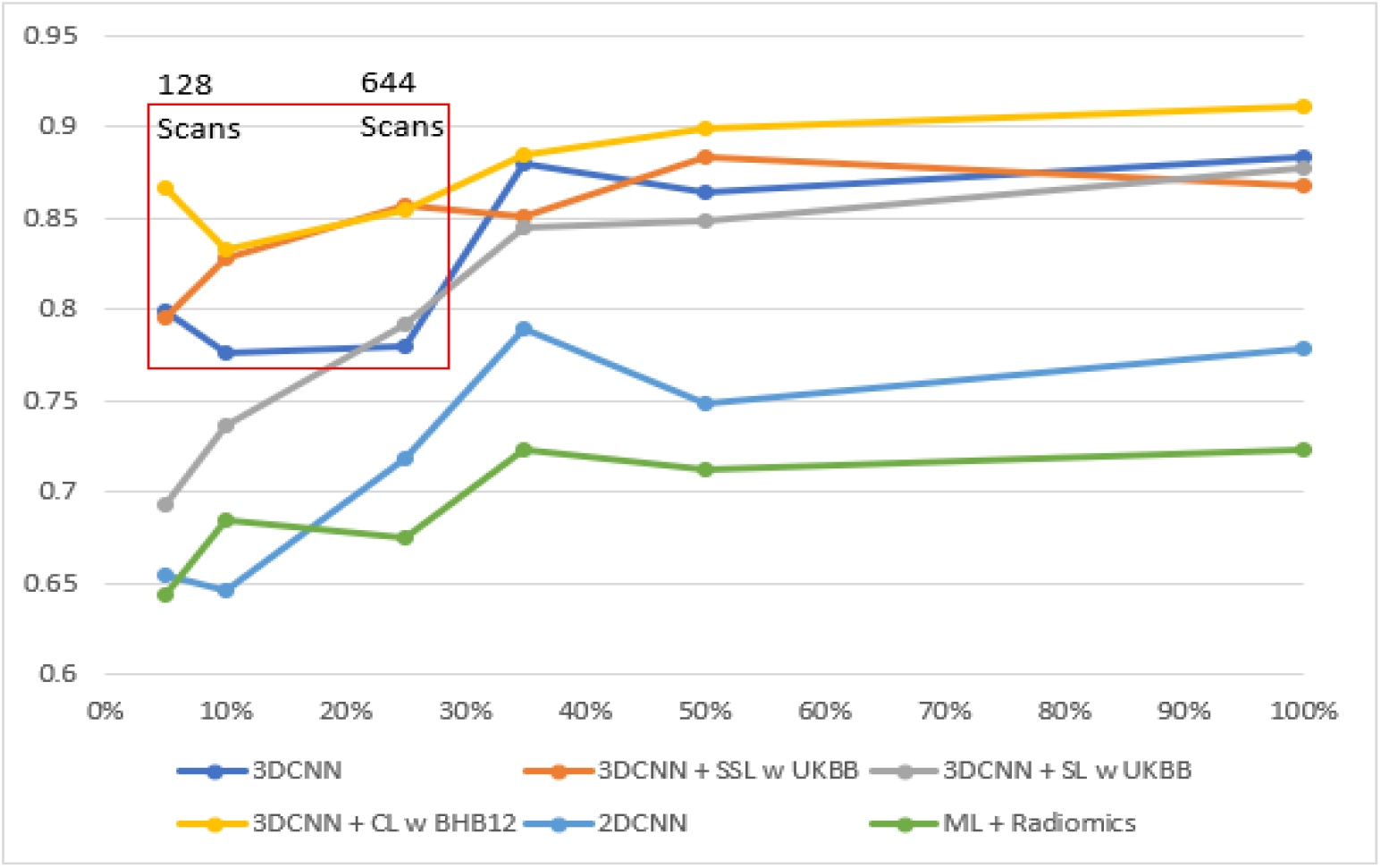
Plot of ADNI test ROC-AUC vs % of training scans, Legend: 5% - 128 scans (121 subjects), 10% - 257 scans (220 subjects), 25% - 644 scans (438 subjects), 35% - 901 scans (525 subjects), 50% - 1,288 scans (628 subjects), 100% - 2,577 scans (747 subjects). The red window in the plot is a representation of a typical low data regime (128 to 644 scans). In our experiments pre-training provided a boost in this limited data region for AD detection relative to when trained from scratch.

The validation loss curves for the 3D CNN trained end-to-end from scratch (*red*) and using modified contrastive pre-training strategy (*green*) for 100% (*left*) and 5% (*right*) of the ADNI training data, are shown in **Figure 2**.

**Figure 2.**
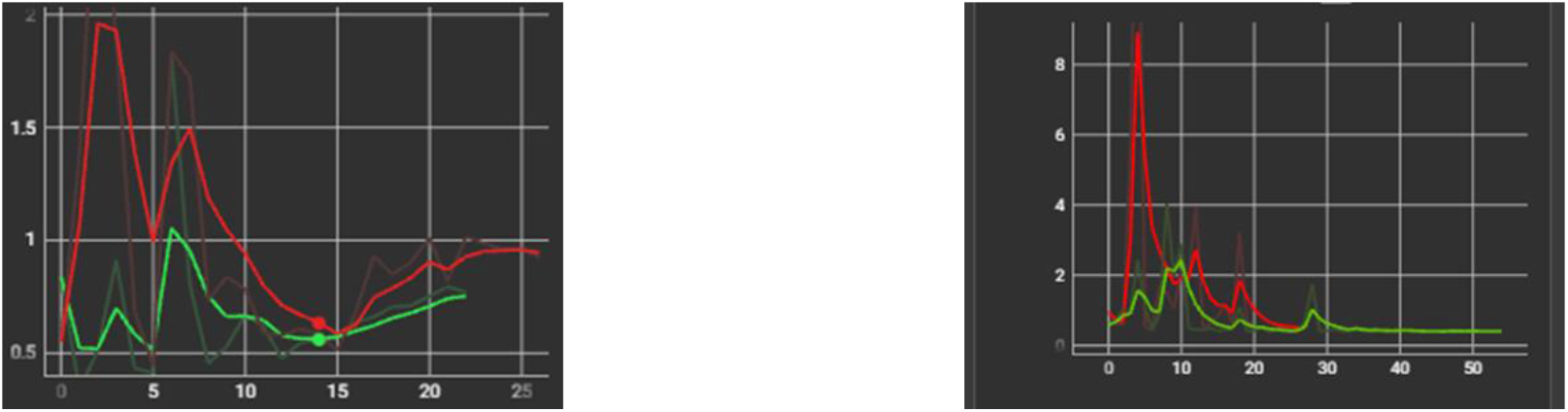
Loss curves during validation. The graphs show Validation loss vs. Number of Epochs, for the 3D CNN trained from scratch (*red*) and 3D CNN + pre-training (*green*). Left: Training on 2,577 scans (747 subjects), Right: Training on 128 scans (121 sub) from ADNI.

The models presented in **Table 3** were further evaluated on an out-of-distribution OASIS dataset; results are summarized in **Table 4**.

**Table 3.** This table extends the test results in **Table 1** with other performance metrics on the ADNI test set for the low training data regimes (10% and 25% training data), where the benefit of pre-training is further highlighted.

| ADNI Test | 25% Training Data |  |  |  | 10% Training Data |  |  |  |
| --- | --- | --- | --- | --- | --- | --- | --- | --- |
|  | Accuracy | F1 | Precision | Recall | Accuracy | F1 | Precision | Recall |
| **3D CNN** | 0.737<br>(0.086) | 0.670<br>(0.122) | 0.702<br>(0.098) | 0.644<br>(0.141) | 0.739<br>(0.016) | 0.663<br>(0.025) | 0.737<br>(0.018) | 0.602<br>(0.030) |
| **3D CNN +**<br>**SSL w**<br>**UKBB** | 0.799<br>(0.032) | 0.746<br>(0.052) | 0.801<br>(0.009) | 0.703<br>(0.089) | 0.783<br>(0.004) | 0.732<br>(0.006) | 0.770<br>(0.022) | 0.700<br>(0.027) |
| **3D CNN +**<br>**SL w UKBB** | 0.719<br>(0.016) | 0.665<br>(0.011) | 0.683<br>(0.058) | 0.659<br>(0.067) | 0.673<br>(0.005) | 0.668<br>(0.015) | 0.593<br>(0.012) | 0.749<br>(0.032) |
| **3D CNN +**<br>**CL w**<br>**BHB12** | 0.804<br>(0.009) | 0.774<br>(0.004) | 0.761<br>(0.022) | 0.788<br>(0.015) | 0.773<br>(0.023) | 0.721<br>(0.029) | 0.755<br>(0.030) | 0.691<br>(0.033) |
| **2D CNN** | 0.652<br>(0.028) | 0.645<br>(0.022) | 0.573<br>(0.030) | 0.743<br>(0.046) | 0.633<br>(0.010) | 0.523<br>(0.043) | 0.585<br>(0.012) | 0.478<br>(0.075) |
| **2D CNN +**<br>**ImageNet** | 0.783<br>(0.034) | 0.746<br>(0.035) | 0.746<br>(0.048) | 0.746<br>(0.021) | 0.732<br>(0.055) | 0.709<br>(0.031) | 0.679<br>(0.087) | 0.755<br>(0.043) |
| **RF +**<br>**radiomics** | 0.648<br>(0.019) | 0.569<br>(0.049) | 0.593<br>(0.023) | 0.554<br>(0.087) | 0.661<br>(0.006) | 0.587<br>(0.008) | 0.609<br>(0.016) | 0.568<br>(0.030) |

**Table 4.** Summary of the various model architectures we trained with different pre-training strategies evaluated on the out-of-distribution OASIS3 dataset, which was collected across several ongoing projects at the Washington University, St. Louis, Alzheimer’s Disease Research Center, over the course of 30 years. This table captures the ability of the various models+pre-training strategies to generalize on an independent dataset from a different imaging site that it has not been trained on. *The pre-training performed originally in ^13^ included OASIS3 on an unrelated task, so we note that there is an overlap with our test set for this model alone (albeit not on the same task).

| OASIS3 T1-w Test ROC AUC | 5% | 10% | 25% | 35% | 50% | 100% |
| --- | --- | --- | --- | --- | --- | --- |
| **3DCNN** | 0.792 (0.005) | 0.815 (0.010) | 0.796 (0.035) | 0.808 (0.007) | 0.815 (0.012) | 0.829 (0.004) |
| **3DCNN + SSL w UKBB** | 0.805 (0.007) | 0.796 (0.014) | 0.813 (0.015) | 0.798 (0.042) | 0.813 (0.021) | 0.819 (0.027) |
| **3DCNN + SL w UKBB** | 0.745 (0.002) | 0.786 (0.003) | 0.813 (0.007) | 0.822 (0.008) | 0.830 (0.005) | 0.844 (0.002) |
| **3DCNN + CL w BHB12** | 0.837 (0.008) | 0.813 (0.021) | 0.824 (0.011) | 0.828 (0.003) | 0.811 (0.011) | 0.812 (0.007) |
| **2DCNN** | 0.632 (0.087) | 0.648 (0.031) | 0.751 (0.029) | 0.764 (0.021) | 0.786 (0.019) | 0.788 (0.009) |
| **2DCNN + ImageNet** | 0.792 (0.022) | 0.810 (0.006) | 0.824 (0.009) | 0.590 (0.179) | 0.656 (0.138) | 0.583 (0.168) |
| **RF + radiomics** | 0.702 (0.002) | 0.725 (0.009) | 0.722 (0.036) | 0.755 (0.002) | 0.746 (0.004) | 0.740 (0.003) |

**Figure 3.**
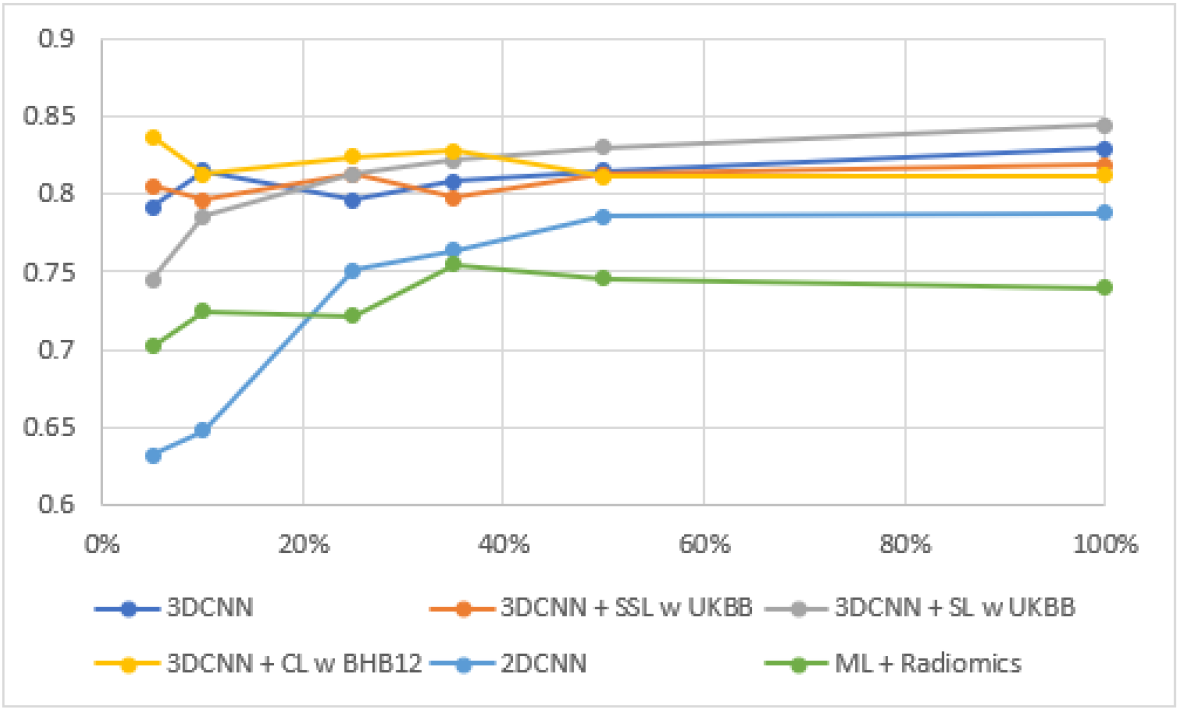
Plot of OASIS3 test ROC-AUC vs % of the available scans used for training. The color key shows how different methods perform after training them using 5% - 128 of the available scans (121 subjects), 10% - 257 scans (220 subjects), 25% - 644 scans (438 subjects), 35% - 901 scans (525 subjects), 50% - 1,288 scans (628 subjects), 100% - 2,577 scans (747 subjects). The 3D CNN with supervised learning performs the best, given enough training data, while the pre-training of models with contrastive learning yields reasonably robust performance (over 0.8) for all training set sizes examined here. Curves are not monotonically increasing due to the random sampling of the training sets and the limited number of runs averaged here.

In **Figure 4**, the UMAP plots visualize the embedding space of the 3D CNN model on the ADNI test set. The two classes (AD/CN) are color coded, as shown in the plots’ legend. The left half of the figure represents the model trained from scratch and the right half represents the model fine-tuned after pre-training on all of the ADNI training data.

**Figure 4.**
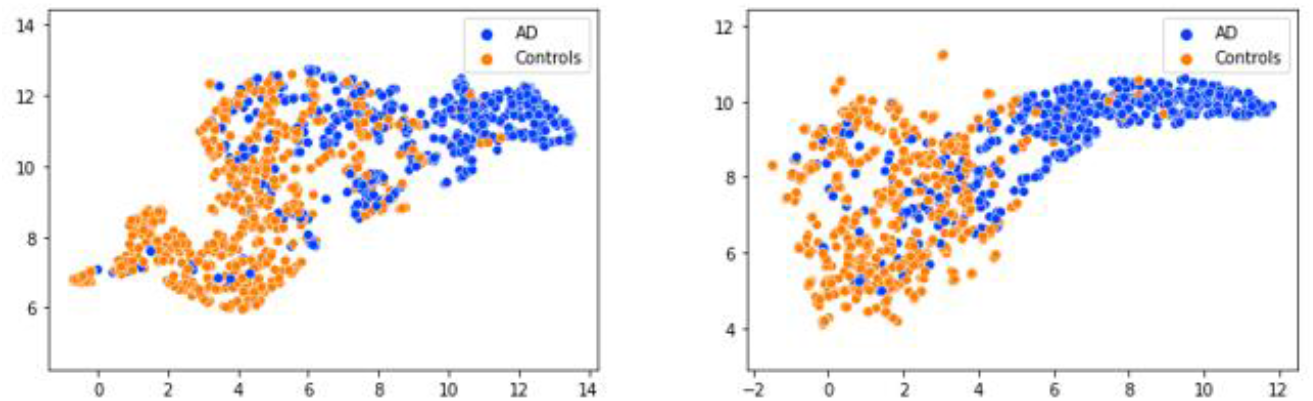
UMAP visualization of the 3D CNN’s embeddings for the two classes: AD and Controls from the ADNI test set. Left: 3D CNN without pre-training, Right: 3D CNN with pre-training.

**Figure 5** shows saliency maps superimposed on the subject’s MRI scans when tested with the 3D CNN trained from scratch (*left*) and the 3D CNN with modified contrastive pre-training (*right*). The three orientations i.e., axial, coronal and sagittal, are represented from left to right in this figure.

**Figure 5.**
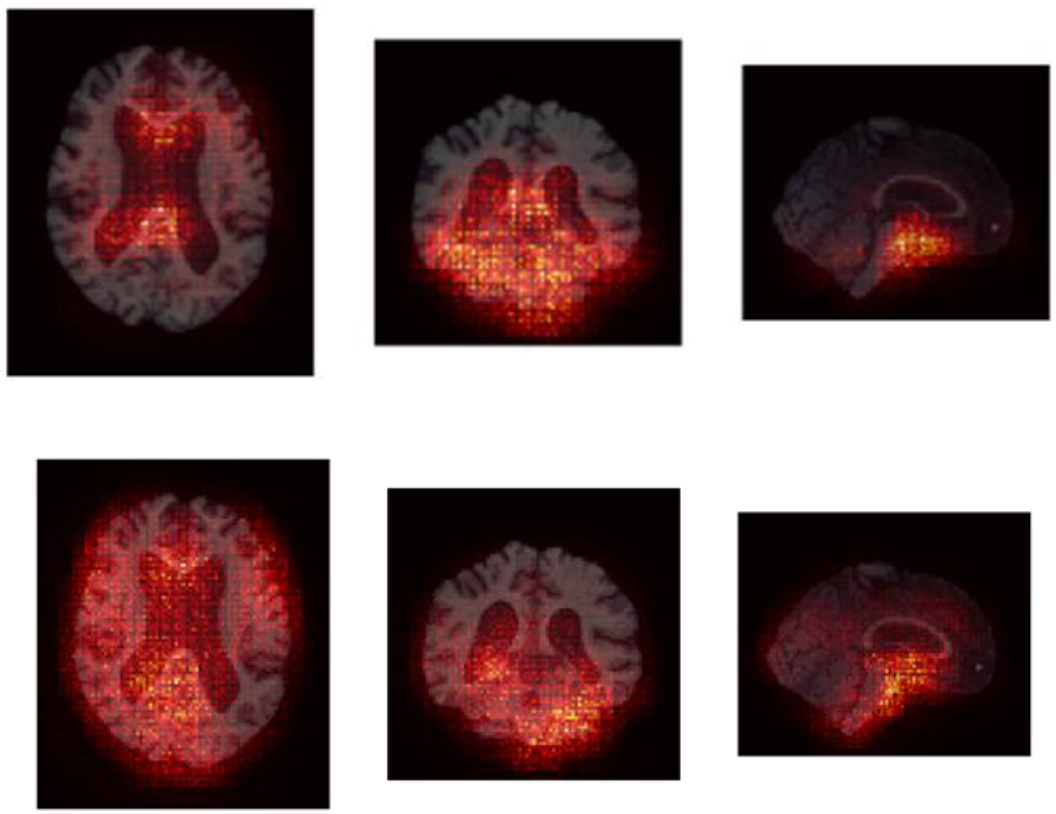
Saliency map visualizations of the models’ predictions for specific axial, coronal and sagittal brain sections. Maps are shown as an average of 10 test subjects from ADNI. Top: 3D CNN without pre-training, Bottom: 3D CNN with pre-training. Additional activation areas for AD Detection in the image are identified by the pre-trained models.

## 5. DISCUSSION AND FUTURE WORK

In this work, we presented results comparing pre-training strategies for Alzheimer’s disease detection from brain MRI. The methods included supervised within-MRI domain, supervised out-of-domain, self-supervised learning, and contrastive learning, using different architectures with T1-w scans. We validated the models on the ADNI test set and an independent out-of-distribution cohort from OASIS3.

A key question about pre-training is whether it boosts tasks performance in general, and especially when the amount of training data for the new task is limited. Even with the full training set of over 2,000 scans, the radiomics methods performed relatively poorly, while the pre-trained models performed much better. Upon fine-tuning all the layers of the 3D CNN (pre-trained in a supervised manner on the sex classification task) for the AD classification task with 25% of the training data, performance of the 3D CNN improved on the ADNI test set with an increase in the average ROC-AUC by 1.2% (0.670 to 0.746), and an increase in the recall of 1.5% (0.644 to 0.659). Self-supervised pre-training improved performance with an increase in the average ROC-AUC by 7.7% (0.780 to 0.857), and an increase in the F-1 score of 7.6% (0.670 to 0.746). The contrastive learning-based pre-training improved performance most of all - with an increase in the average ROC-AUC by 7.4% (0.780 to 0.854), and an increase in the F-1 score of 10.4% (0.670 to 0.774). Pre-training was also effective in experiments that used very limited amounts of training data for the AD prediction task. Fine-tuning the 3D CNN with only 10% of the training data (only 257 MRI scans) using self-supervised pre-training improved performance with an increase in the average ROC-AUC by 5.2% (0.776 to 0.828), and an increase in the F-1 score of 6.9% (0.663 to 0.732). The contrastive learning-based pre-training also improved performance - with an increase in the average ROC-AUC of 5.7% (0.776 to 0.833), and an increase in the F-1 score of 5.8% (0.663 to 0.721).

From our experiments, the 3D CNN T1-w scans benefited the most in low data regimes, i.e., when using only 10% to 25% of the total training data. The validation loss curve in **Figure 2** shows relatively lower losses over time for the pre-trained model, and the model is also more stable. The UMAP visualization is a projection of the high-dimensional embedding space of the trained model. Our UMAP in **Figure 4** shows the subjects in the AD/control classes from the ADNI test set. Qualitatively, the UMAP clustering of the classes is improved and appears more compact in the pre-trained case with fewer outliers, as seen in **Figure 4**. The saliency maps in **Figure 5** are based on the gradients of the predicted class with respect to the input pixels; they highlight regions of the subject’s brain scan that contribute maximally to the final prediction score. The maps for the model with pre-training (bottom right) cast attention on more brain regions, relative to the model trained from scratch. It is not clear *a priori* that this is better, but the maps offer evidence that informative features have been found throughout the image. Some of the key considerations in our pre-training work include the domains of the pre-training and downstream tasks, model size and architecture, number of layers being fine-tuned, effect and state of the batch normalization layers during fine-tuning and tuning of the hyperparameters. We noticed that fine-tuning all the layers of the 2D CNN pre-trained on ImageNet did not significantly improve performance of the baseline trained from scratch, but it did seem to have a positive effect when using less data to fine-tune, as seen in **Tables 2 and 3**. Although there have been papers reporting performances with very high AUCs, exceeding 0.95, Wen et al.^6^ note the possibility of data leakage in many published models for AD classification. Ultimately, the inter-rater variability of the radiologist could be a reasonable target for machine learning models to attain.

This study has several limitations, which we will address in future work. First, AD classification often uses other data sources, such as amyloid- or tau-sensitive PET, FDG-PET, and also clinical data such as the MMSE and other tests of dementia severity. Blood markers are also showing promise for AD classification. An MRI-based model is unlikely to be used in isolation, without these additional data sources, but for simpler benchmarking we did not include them here, to better understand how performance depends on the training approaches used. Second, in AD clinical research, the focus has somewhat evolved from binary AD classification to subtyping and staging, ^5 30^ as well as future prognosis. These tasks presumably may also benefit from the pre-training methods examined here and are a topic for future work. Finally, very recent work has pre-trained MRI deep learning methods on YouTube videos,^31^ raising the intriguing possibility that natural image or video pre-training of MRI deep learning models may perform as well, in some cases, as pre-training on tasks in the medical domain.

In our immediate future work, we plan to also conduct experiments in a federated setting to incorporate data from multiple sites in a secure manner for dementia classification, subtyping, and prognosis. With additional data - including longitudinal scans - the models could be used to predict progression of dementia or classification of other neurological diseases with limited publicly available data such as Parkinson’s disease.^4^

## 6. CONCLUSIONS

In this work, we show the benefits of various pre-training strategies to improve automated detection of Alzheimer’s disease based on brain MRI. Our visualizations provide additional interpretation of the models’ predictions and how different parts of the brain image contribute to the output score. Model pre-training could also accelerate the application of deep learning models to other diseases where publicly available neuroimaging data is limited.

## Acknowledgments

This work was supported by the U.S. National Institutes of Health, under NIH grant U01 AG068057, and by DARPA under Agreement HR00112090104.

